# Neuronal primary cilia are not required for hippocampal circuit function or behavior in adult mice

**DOI:** 10.64898/2026.08.24.746770

**Authors:** Tae-Yeon Eom, Ildar T. Bayazitov, Brett J.W. Teubner, Donnie Eddins, Stanislav S. Zakharenko

**Affiliations:** Division of Neural Circuits and Behavior, Department of Developmental Neurobiology, St. Jude Children’s Research Hospital, Memphis, TN 38105, USA

**Keywords:** Primary cilia, Ift88, Neuronal activity, Hippocampus, Learning and memory

## Abstract

Primary cilia, which are present in most brain cells, are essential for brain development and function. During early brain development, dysfunction of the primary cilia can lead to a broad spectrum of disorders, collectively termed ciliopathies, that include brain malformations and intellectual disability. Although the role of primary cilia in brain development is well-established, cilia-mediated signaling in mature neurons and the contribution of cilia to neuronal circuit function remain controversial. Using mouse genetic and behavioral studies, single-cell electrophysiology, and 2-photon imaging, we show that deletion of primary cilia from adult hippocampal neurons is not required for hippocampal circuit function or behavior. Chronic genetic deletion or acute laser ablation of primary cilia from mature pyramidal neurons in the CA1 or CA3 regions of the hippocampus did not affect neuronal excitability, basal synaptic transmission, or long-term synaptic plasticity at excitatory CA3–CA1 hippocampal synapses. Moreover, the loss of primary cilia did not affect hippocampal-dependent learning and memory or anxiety-like behaviors. These results challenge the prevailing view of cilia function in mature hippocampal neurons and suggest that neuronal cilia in the adult hippocampus do not serve as major signaling hubs for pathways essential for neuronal function or behavior.

## Introduction

Primary cilia are solitary, nonmotile, microtubule-based cellular projections that extend from the surface of most vertebrate cells into the extracellular space. Although primary cilia were long-considered vestigial remnants, recent findings have revealed their fundamental roles in brain development and homeostasis, including regulation of cell fate determination, migration, and differentiation, through their function as a signaling hub^1–5^. Cilia are assembled by the highly conserved process of intraflagellar transport (IFT) and serve as sensory antennae that receive and transduce signals from adjacent cells or the surrounding extracellular environment^6–8^. Cilia dysfunction can lead to a broad spectrum of disorders collectively termed “ciliopathies,” which include brain malformations, kidney disease, vision impairment, intellectual disability, and obesity^9–11^. Furthermore, disruption of cilia formation results in various neural phenotypes, such as cerebellar malformations, abnormal neuronal migration, and defects in adult neurogenesis^12–15^. Recent studies have also shown a link between cilia and neuronal activity, as well as behavior, in mouse models^16–20^. Although evidence supporting the role of primary cilia in neuronal activity and behavior is emerging, these effects may be secondary to alterations in other intracellular signaling pathways or changes in cilia of immature neuronal progenitors, rather than reflecting the direct role of the cilia in mature neurons. To directly test whether cilia contribute to the function of mature neurons, neural circuits, and the behaviors they control, we deleted primary cilia later during postnatal development by using genetic or laser-ablation tools.

## Results

### Genetic ablation of primary cilia in postmitotic hippocampal neurons

To examine the functional role of primary cilia in the adult brain, we conditionally deleted primary cilia from postmitotic hippocampal neurons of mice carrying a floxed allele of the *Ift88* gene (*Ift88^fl/fl^*mice), which is essential for cilia formation and function^21^. To determine whether cilia have a role in synaptic transmission and plasticity that is separate from their developmental effect, we generated *Ift88–*conditional knockout (cKO) mice in which the deletion of *Ift88* was not only restricted to the forebrain pyramidal neurons but also restricted temporally to postnatal development; thus, *Ift88* deletion became apparent only after postnatal day (P)14, with the proportion of cells showing deletion increasing with age^22–25^. In these experiments, we focused on the excitatory synapses between CA3 and CA1 pyramidal neurons (CA3–CA1 synapses) of the hippocampus, because synaptic transmission and long-term synaptic plasticity at these synapses are well-characterized and have been strongly linked to hippocampus-dependent behaviors^26–28^.

We also crossed *Ift88^fl/fl^* mice with mice expressing Cre recombinase (Cre) under control of calcium/calmodulin-dependent protein kinase II alpha promoter (*Camk2a^Cre^* mice) to delete the *Ift88* gene in postmitotic neurons in the forebrain, including CA1 neurons^22^. Cre expression in these mice begins at approximately P18, with recombination fully established by 4 weeks of age^22^. To delete the *Ift88* gene in CA3 neurons, we crossed *Ift88^fl/fl^* mice with mice expressing Cre under control of glutamate receptor, ionotropic, kainite 4 promoter (*Grik4*^Cre^ mice)^29^. Cre expression begins in these mice at approximately P14, with recombination fully established by 8 weeks of age^29^.

To verify Cre expression, we crossed each Cre line with the Ai14 reporter strain (*ROSA26-CAG-Stop^fl/fl^-tdTomato*). We used 3- to 5-month-old mice of both sexes. Robust tdTomato expression was detected throughout the hippocampus (CA1-3 and dentate gyrus [DG]) in *Camk2a^Cre^;Ai14* mice and in the CA3 neurons and DG of *Grik4^Cre^;Ai14* mice (**Fig. 1a, d**). To confirm the ablation of primary cilia in these lines, we co-labeled cells with tdTomato and the ciliary marker adenylate cyclase III (ACIII). This approach showed a marked depletion of cilia in CA1 neurons of *Ift88^fl/fl^;Camk2a^Cre^;Ai14* mice and in CA3 neurons of *Ift88^fl/fl^;Grik4^Cre^;Ai14* mice compared with control littermates (**Fig. 1b, e**). We also examined cilia presence in hippocampal subfields (**Supplementary Figs. S1, S2**) and quantified the extent of cilia loss in each region. In Cre-expressing neurons, we observed a significant reduction in the number of ACIII^+^ cilia in the CA1 (97%), CA2 (81%), and DG (45%) neurons of *Ift88^fl/fl^;Camk2a^Cre^;Ai14* mice and in the CA3 neurons (90%) of *Ift88^fl/fl^;Grik4^Cre^;Ai14* mice, relative to matched controls (**Fig. 1c, f**). These findings reflect the differential Cre expression and recombination efficiency across hippocampal subregions. To assess whether cilia depletion affected hippocampal neurogenesis, we quantified doublecortin (DCX)^+^ cells, a marker of young neurons, in the granule cell layer of the DG. We observed no significant differences between genotypes (**Supplementary Fig. S3**). Together, these results indicate that primary cilia are efficiently and selectively ablated in mature hippocampal neurons by using the *Ift88* floxed allele, in combination with *Camk2a^Cre^* or *Grik4^Cre^* drivers, thus providing a spatially and temporally controlled model of cilia loss in the adult hippocampus.

**Fig. 1.**
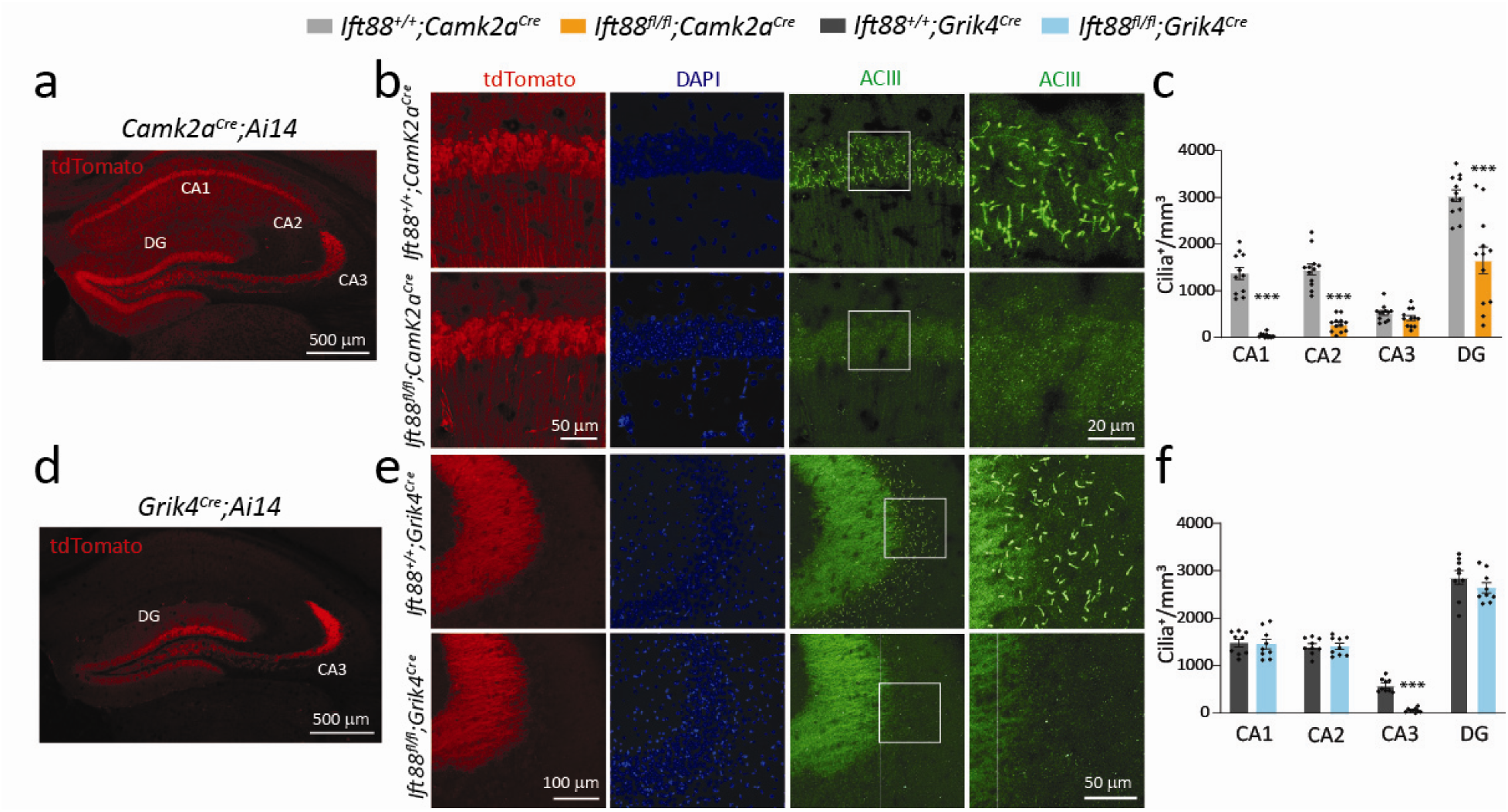
Conditional deletion of primary cilia in mature hippocampal neurons. (**a**) A representative image of a tdTomato^+^ (red) hippocampal section from a *Camk2a^Cre^;Ai14* mouse. (**b**) Images of a cilia marker ACIII^+^ (green) and tdTomato^+^ (red) neurons in the CA1 region of *Ift88^+/+^;Camk2a^Cre^;Ai14* mice and *Ift88^fl/fl^;Camk2a^Cre^;Ai14* mice. The boxed region is shown at a higher magnification to better visualize primary cilia. Nuclei were counterstained with DAPI (blue). (**c**) Quantifications of primary cilia (quantified as ACIII^+^ organelles) in CA1-3 and dentate gyrus (DG) of control (*Ift88^+/+^;Camk2a^Cre^*; n=4 mice, 12 sections) and mutant (*Ift88^fl/fl^;Camk2a^Cre^*; n=4 mice, 12 sections) mice. Two-way ANOVA (Holm-Sidak’s post hoc): *F*_3, 87_= 10.75, \*\*\**p* <0.0001. (**d**) A representative image of a tdTomato^+^ hippocampal section from a *Grik4^Cre^;Ai14* mouse. (**e**) Images of ACIII^+^ (green) and tdTomato^+^ (red) neurons in the CA3 region of *Ift88^+/+^;Girk4^Cre^;Ai14* mice and *Ift88^fl/fl^;Grik4^Cre^;Ai14* mice. The boxed region is shown at a higher magnification to better visualize primary cilia. Nuclei were counterstained with DAPI. (**f**) Quantifications of primary cilia in hippocampal subregions of control (*Ift88^+/+^;Grik4^Cre^*; n=3 mice, 9 sections) and mutant (*Ift88^fl/fl^;Grik4^Cre^*; n=3 mice, 9 sections) mice. Two-way ANOVA (Holm-Sidak’s post hoc): *F*_3, 64_= 3.93, \*\*\**p*=0.01. Data are presented as mean ± SEM in **c** and **f**.

### Chronic loss of primary cilia in adult mice does not compromise neuronal excitability, synaptic transmission, or short- and long-term synaptic plasticity

We next examined the neurophysiological consequences of primary cilia deletion by whole-cell patch-clamp recordings in acute hippocampal slices from *Ift88*-cKO mice. Deletion of cilia from CA1 or CA3 pyramidal neurons of *Ift88^fl/fl^;Camk2a^Cre^* or *Ift88^fl/fl^;Grik4^Cre^*mice, respectively, did not alter the resting membrane potential (MP; **Fig. 2a, g**), input resistance (**Fig. 2b, h**), rheobase (**Fig. 2c, i**), action potential (AP) firing (**Fig. 2d, j**), AP amplitude (**Fig. 2e, k**), or AP half-width (**Fig. 2f, l**), indicating that neuronal intrinsic excitability was preserved in the absence of primary cilia.

**Fig. 2.**
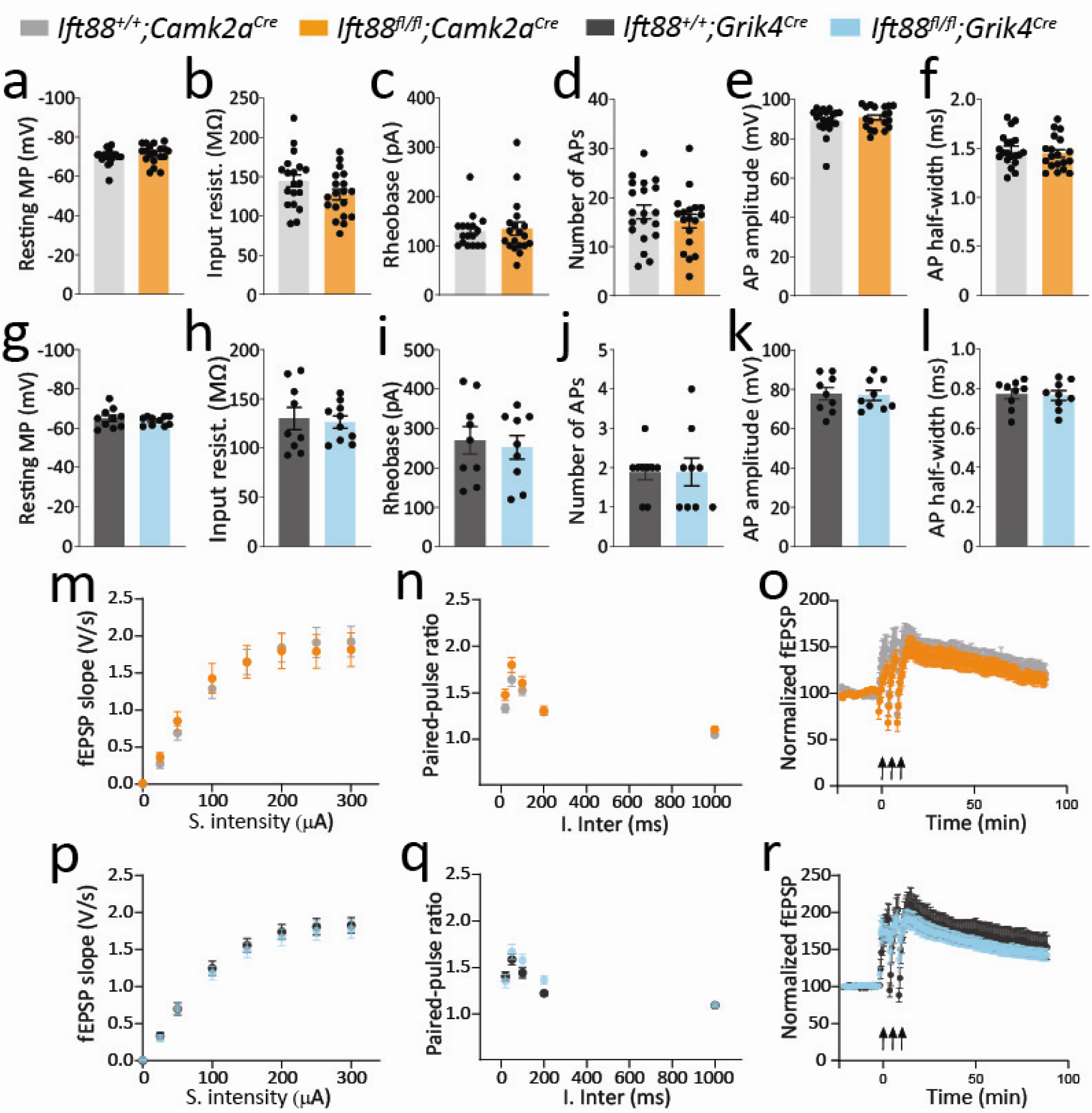
Loss of primary cilia does not alter hippocampal neuronal excitability, synaptic transmission, or synaptic plasticity. Whole-cell recordings from CA1 (**a**-**f**) or CA3 (**g**-**l**) pyramidal neurons showed no differences in resting membrane potential (MP; **a**, **g**), input resistance (**b**, **h**), rheobase (**c**, **i**), number of action potentials (APs; **d**, **j**), AP amplitude (**e**, **k**), or AP half-width (**f**, **l**) between control (*Ift88^+/+^;Camk2a^Cre^*, n=19 cells from 4 mice; *Ift88^+/+^;Grik4^Cre^*, n=10 cells from 4 mice) and mutant (*Ift88^fl/fl^;Camk2a^Cre^*, n=20 cells from 5 mice; *Ift88^fl/fl^;Grik4^Cre^*, n=10 cells from 4 mice) mice. Welch’s unpaired *t*-test: (**a**) *p*=0.368; (**b**) *p*=0.1004; (**c**) *p*=0.757; (**d**) *p*=0.351; (**e**) *p*=0.375; (**f**) *p*=0.513; (**g**) *p*=0.444; (**h**) *p*=0.77; (**i**) *p*=0.706; (**j**) *p* >0.999; (**k**) *p*=0.868; (**l**) *p*=0.823. (**m**-**r**) Primary cilia deletion also did not affect basal synaptic transmission, measured by the input/output relationship (field excitatory postsynaptic potential [fEPSP] slope; (**m**, **p)**, paired-pulse ratio (**n**, **q**), or long-term potentiation (**o**, **r**) in *Ift88^+/+^;Camk2a^Cre^* (3 mice, 13 slices), *Ift88^fl/fl^;Camk2a^Cre^* (4 mice, 16 slices), *Ift88^+/+^;Grik4^Cre^* (4 mice, 21 slices), and *Ift88^fl/fl^;Grik4^Cre^* (4 mice, 21 slices). The three arrows (in **o** and **r**) indicate induction of long-term potentiation. Two-way repeated-measures ANOVA: in (**m**) *p*=0.94, *F*_1, 27_=0.006, (**n**) *p*=0.16, *F*_1, 27_=2.129, (**o**) *p*=0.45, *F*_1, 27_=0.58, (**p**) *p*=0.69, F_1, 40_=0.16, (**q**) *p*=0.29, *F*_1, 40_=1.13, and (**r**) *p*=0.61, *F*_1, 40_=0.268. Abbreviations: I. inter., interpulse interval; S. stimulation. Data are presented as mean ± SEM.

Similarly, using field recordings in acute hippocampal slices, we determined that cilia deletion did not affect basal synaptic transmission (**Fig. 2m, p**), short-term plasticity measured by paired-pulse ratio (**Fig. 2n, q**), or long-term potentiation (LTP) at CA3–CA1 synapses induced by tetanization of Schaffer collaterals (**Fig. 2o, r**) in either *Ift88^fl/fl^;Camk2a^Cre^*or *Ift88^fl/fl^;Grik4^Cre^* mice, compared with their respective control littermates. Together, these data indicate that disruption of primary cilia in hippocampal pyramidal neurons does not impair intrinsic excitability, basal synaptic transmission, or synaptic plasticity.

### Acute or viral-mediated deletion of cilia from adult hippocampal neurons does not affect neuronal excitability or synaptic function

The lack of effect of chronic deletion of primary cilia on neuronal or synaptic function in *Ift88^fl/fl^;Camk2a^Cre^* or *Ift88^fl/fl^;Grik4^Cre^* mice could be explained by compensatory mechanisms. To rule out this possibility, we performed acute or viral-mediated ablation of primary cilia in hippocampal pyramidal neurons. To acutely ablate the primary cilia, we performed whole-cell patch-clamp recordings of pyramidal hippocampal neurons, while filling each neuron with the red fluorescent dye Alexa Fluor 594 to visualize its morphology. We performed these experiments in acute hippocampal slices from *Arl13b-GFP* mice^30^, which enabled us to visualize the primary cilia by green fluorescence protein (GFP) via 2-photon imaging. To ablate the primary cilium in the recorded neuron, we used a short, targeted 2-photon laser exposure aimed at the proximal end of the cilium. This exposure effectively removed the cilium without affecting the recorded neuron (**Fig. 3a**, **b**). Indeed, current-clamp recordings showed that acute cilium ablation did not alter intrinsic neuronal excitability, as measured by resting MP, input resistance, rheobase, AP firing, and AP half-width (**Fig. 3c**-**f****, h**), though the AP amplitude was reduced a moderate but significant amount (**Fig. 3g**). Firing patterns were similar before (cilia intact) and after (cilia ablated) laser ablation (**Fig. 3i**).

**Fig. 3.**
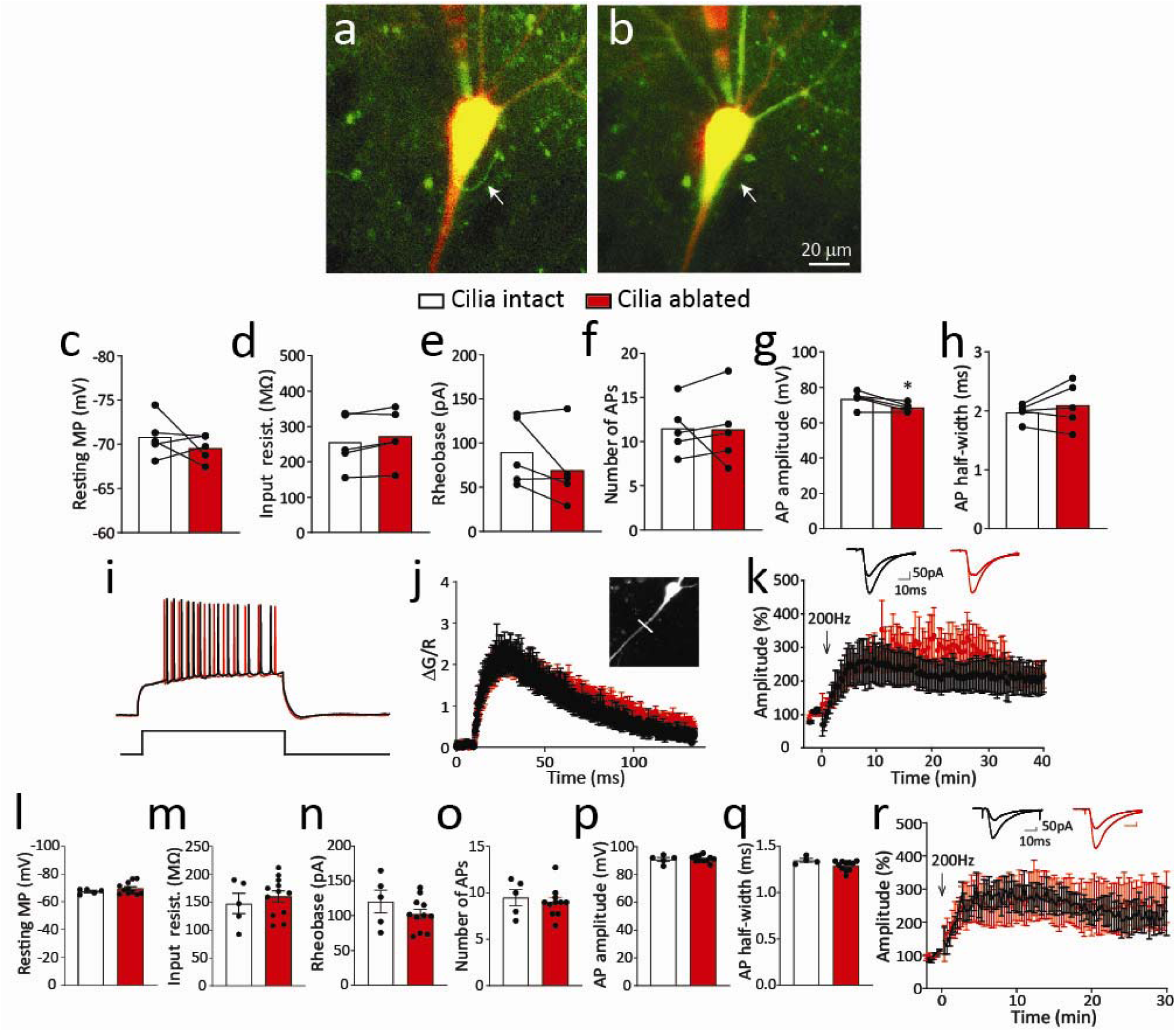
Acute or virus-mediated cilia deletion does not alter passive membrane properties, neuronal excitability, synaptic transmission, or synaptic plasticity in the hippocampus. (**a**) Representative image of a CA1 pyramidal neuron from an *Arl13b-GFP* mouse showing the primary cilium labeled with GFP (green, arrow). The neuron was filled with Alexa Fluor 594 (red) through the recording pipette. (**b**) The same neuron after laser ablation of the primary cilium. (**c**-**h**) Acute cilium ablation did not alter the resting membrane potential (MP; **c**), input resistance (**d**), rheobase (**e**), number of action potentials (APs; **f**), or the AP half-width (**h**), though the AP amplitude was modestly but significantly reduced (**g**). Paired *t*-test: (**c**) *t*_4_=0.946, *p*=0.398; (**d**) *t*_4_=2.564, *p*=0.062; (**e**) *t*_4_=1.649, *p*=0.175; (**f**) *t*_4_=0.073, *p*=0.945; (**g**) *t*_4_=3.121, * *p*=0.036; and (**h**) *t*_4_=1.014, *p*=0.368. (**i**) Representative current-clamp recordings before (black) and after (red) laser ablation of the cilium. (**j**) Dendritic calcium transients evoked by trains of back-propagating APs were unchanged after cilium ablation. The inset shows the apical dendritic region used for calcium measurements (ΔG/R). Paired *t*-test: *t*_4_=1.236, *p*=0.284. (**k**) LTP in CA1 was unaffected by acute presynaptic cilium ablation. Insets show representative EPSCs before and after ablation. Two-tailed Student’s *t*-test: *t*_9_=0.978, *p*=0.353. (**l**-**r**) AAV-GFP-Cre–mediated cilia deletion in CA1 neurons of *Ift88^fl/fl^* mice did not alter resting MP (**l**), input resistance (**m**), rheobase (**n**), AP number (**o**), AP amplitude (**p**), or AP half-width (**q**). Unpaired *t*-test: (**l**) *t*_14_=1.075, *p*=0.30; (**m**) *t*_15_=0.661, *p*=0.519; (**n**) *t*_14_=1.201, *p*=0.25; (**o**) *t*_14_=0.55, *p*=0.591; (**p**) *t*_14_=0.365, *p*=0.721; (**q**) *t*_13_=1.986, *p*=0.069. (**r**) Virus-mediated cilia deletion had no effect on LTP in CA1 neurons. Insets show representative EPSCs before and after ablation. Welch’s unpaired *t*-test: *p*= 0.582. Data are presented as mean ± SEM.

To test if acute cilia removal affected dendritic signaling, we filled a neuron with the calcium fluorescent indicator Fluo-5F (green) in addition to Alexa Fluor 594 (red)^31,32^. We then measured dendritic calcium transients induced by a train of back-propagating APs (20 APs at 100 Hz), before and after cilia ablation in the same neurons. Calcium transients, which were measured as Fluo-5F fluorescence normalized to Alexa Fluor 594 fluorescence, were unchanged after cilium ablation (**Fig. 3j**).

Using voltage-clamp recordings, we then tested if acute ablation of the primary cilia affected long-term plasticity in acute hippocampal slices. In these experiments, we compared CA1 neurons with intact primary cilium and CA1 neurons with the primary cilium ablated 5–10 min prior to LTP induction by Schaffer collateral tetanization. These experiments showed that LTP at CA3–CA1 excitatory synapses was also unaffected by acute cilium ablation in CA1 pyramidal neurons (**Fig. 3k**).

We then tested the effect of the primary cilia on synaptic transmission between connected pairs of neurons. Because paired recordings in the hippocampus are technically challenging, we recorded from pairs of well-interconnected layer 4 pyramidal neurons in the auditory cortex^32^ of *Arl13b-GFP* mice. After filling these neurons with Alexa Fluor 594, we visualized red-labelled cell bodies with green-labelled cilia. Injection of depolarizing current in one neuron produced AP firing in current-clamp mode and a time-locked excitatory postsynaptic current (EPSC) recorded in voltage-clamp mode in the second neuron, indicating that the first neuron was synaptically connected to the second neuron. After establishing a 5-min baseline, we removed the cilium from the first neuron by using a 2-photon laser and then continued EPSC recordings for 20–25 min. Consistent with the previous experiment, synaptic transmission between pairs of synaptically connected layer 4 neurons in the auditory cortex remained unchanged after presynaptic cilium ablation (**Supplementary Fig. S4a-c**).

In another set of experiments, we used adeno-associated virus (AAV)-mediated ablation of the primary cilia. We deleted the *Ift88* gene by *in vivo* injecting the AAV encoding Cre-GFP under control of the hSynapsin promoter (AAV-GFP-Cre) into the dorsal hippocampus of *Ift88^fl/fl^*mice, resulting in primary cilia ablation in hippocampal neurons (**Supplementary Fig. S5**). We then compared resting MP, input resistance, rheobase, AP firing, and AP waveforms between GFP^+^ neurons in the CA1 pyramidal neurons of *Ift88^fl/fl^* mice and control *Ift88^+/+^*mice injected with AAV-GFP-Cre. Using this approach, we detected no difference in neuronal excitability between mutant and control neurons (**Fig. 3l-q**).

We used a similar approach to measure long-term synaptic plasticity at CA3–CA1 synapses following virus-mediated cilia ablation in these mice. We recorded from GFP^+^ neurons in voltage-clamp mode and induced LTP by tetanizing Schaffer collaterals. LTP induced in *Ift88^fl/fl^*mice injected with AAV-GFP-Cre was similar in magnitude to that induced in *Ift88^+/+^* mice injected with AAV-GFP-Cre (**Fig. 3r**). Together, these findings indicate that neither acute nor virus-mediated deletion of primary cilia in mature neurons impairs intrinsic neuronal excitability, synaptic transmission, or synaptic plasticity, further supporting the notion that primary cilia are not required for normal neuronal function in the adult hippocampus.

### Primary cilia ablation in mature hippocampal neurons does not impair learning, memory, anxiety-like behavior, social behavior, or motor function

Previous studies have reported deficits in hippocampal-dependent learning and memory^16,20,33^ and object recognition memory^17,33^ after disruption of neuronal primary cilia at various ages. To investigate these behavioral outcomes when the primary cilia deletion is restricted to mature neurons, we performed a battery of behavioral tests assessing learning and memory, anxiety-like behavior, social behavior, and motor function. We first assessed associative learning and memory by using contextual and cued fear-conditioning paradigms. Both *Ift88^fl/fl^;Camk2a^Cre^*mice and *Ift88^fl/fl^;Grik4^Cre^* mice exhibited freezing responses comparable to those of their respective control littermates, indicating intact learning and memory for both contextual and cued fear-conditioning tests (**Fig. 4a, e**). In the novel object recognition test, *Ift88^fl/fl^;Camk2a^Cre^* mice and their littermate controls displayed similar preferences for the novel object (**Fig. 4b**). Although the preference was slightly less pronounced, *Ift88^fl/fl^;Grik4^Cre^*mice also showed a level of novel object discrimination comparable to that of controls (**Fig. 4f**). Spatial learning and memory were examined using the Morris water maze. *Ift88^fl/fl^;Camk2a^Cre^* mice exhibited escape latencies that were indistinguishable from those of controls, as well as similar time spent in the target quadrant during both the probe test conducted 1 h after training and the memory-retention test performed 72 h later (**Fig. 4c, d**). Similar results were obtained in *Ift88^fl/fl^;Grik4^Cre^*mice (**Fig. 4g, h**).

**Fig. 4.**
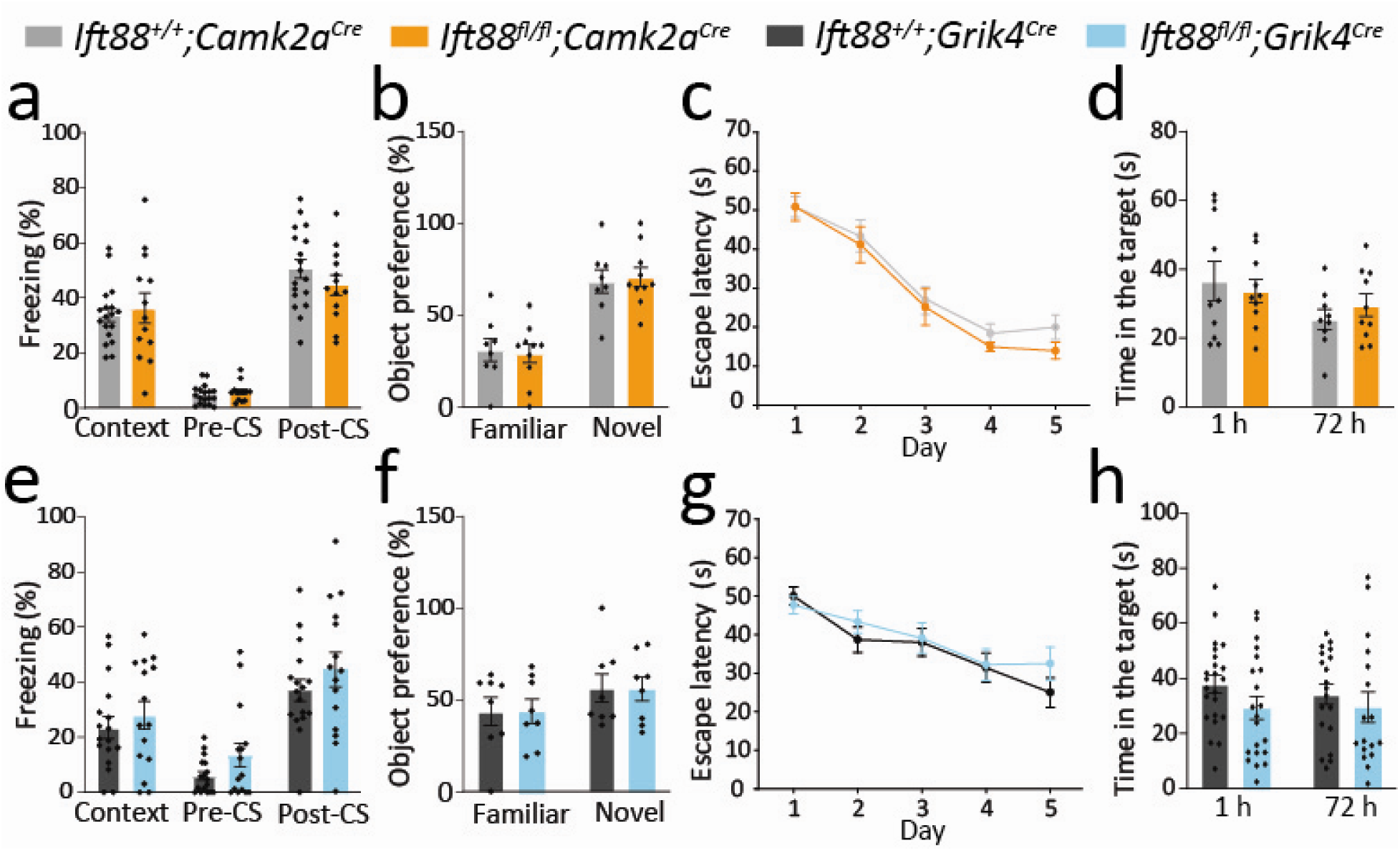
Primary cilia deletion in the postmitotic hippocampal neurons does not affect learning or memory. (**a**, **e**) Fear conditioning showed no differences in freezing during the contextual test (Context), before the conditioned stimulus (Pre-CS), or after the conditioned stimulus (Post-CS) between control (*Ift88^+/+^;Camk2a^Cre^*, n=18 mice; *Ift88^+/+^;Grik4^Cre^*, n=17 mice) and mutant (*Ift88^fl/fl^;Camk2a^Cre^*, n=13 mice; *Ift88^fl/fl^;Grik4^Cre^*, n=15 mice) mice. Two-way ANOVA (Holm-Sidak’s post hoc): (**a**) *F*_2, 87_=1.05, *p*=0.35; (**e**) *F*_2, 90_=0.08, *p*=0.92. (**b**, **f**) Novel object recognition showed no differences in discrimination between familiar and novel objects in control (*Ift88^+/+^;Camk2a^Cre^*, n=8 mice; *Ift88^+/+^;Grik4^Cre^*, n=8 mice) and mutant (*Ift88^fl/fl^;Camk2a^Cre^*, n=10 mice; *Ift88^fl/fl^;Grik4^Cre^*, n=8 mice) mice. Two-way ANOVA (Holm-Sidak’s post hoc): (**b**) *F*_1, 32_=0.12, *p*=0.73; (**f**) *F*_1, 28_=0.005, *p*=0.95. (**c**, **d**, **g**, **h**) Morris water maze test showed no difference in spatial learning during training (escape latency) or memory recall at 1 h or 72 h after training between control (*Ift88^+/+^;Camk2a^Cre^*, n=10 mice, *Ift88^+/+^;Grik4^Cre^*, n=19 mice) and mutant (*Ift88^fl/fl^;Camk2a^Cre^*, n=10 mice, *Ift88^fl/fl^;Grik4^Cre^*; n=17 mice) mice. Two-way ANOVA (Holm-Sidak’s post hoc): (**c**) *F*_4, 90_=0.22, *p*=0.93; (**d**) *F*_1, 35_=0.78, *p*=0.38; (**g**) *F*_4, 170_=0.58, *p*=0.68; (**h**) *F*_1, 76_=0.23, *p*=0.63. Data are presented as mean ± SEM.

To determine whether neuronal cilia ablation affected other behavioral domains, we evaluated anxiety-like behavior, social behavior, and motor function. In the open-field test, time spent in the center zone was comparable between mutant mice and control mice (**Supplementary Fig. S6a, e**). Likewise, the three-chamber social interaction and social recognition test revealed no differences between genotypes (**Supplementary Fig. S6b, c, f, g**), and performance on the rotarod test was also comparable between groups (**Supplementary Fig. S6d, h**). Together, these results indicate that the loss of primary cilia from mature hippocampal neurons does not impair hippocampal-dependent learning and memory, anxiety-like behavior, social behavior, or motor function, further suggesting that neuronal primary cilia are dispensable for these behavioral functions in the adult brain.

## Discussion

Although primary cilia are retained throughout the lifespan of most cells, their contribution to adult brain function through cilia-mediated signaling remains poorly understood, compared with their well-established roles in early brain development^1,4,34^. Moreover, ciliopathies encompass a wide spectrum of disorders, ranging from developmental abnormalities to neurodegenerative diseases that occur at different stages of development^10,11^. However, the role of primary cilia in neuronal and circuit function in the adult brain and whether ciliary dysfunction contributes to these conditions remains unclear. Unexpectedly, we found that ablation of neuronal cilia in the adult mouse brain did not impair neuronal function or behavior. Conditional deletion of primary cilia in mature hippocampal neurons did not affect intrinsic excitability, synaptic transmission, or short- or long-term synaptic plasticity. Furthermore, these mice exhibited no deficits in hippocampal-dependent learning and memory, anxiety-like behavior, social behavior, or motor function. These findings refine the current understanding of primary cilia function in mature hippocampal neurons and suggest that, unlike during development, in the adult hippocampus neuronal cilia are not major signaling hubs required for maintaining neuronal function or behavioral performance.

Neuronal cilia dynamically house various signaling molecules, including G protein–coupled receptors, receptor tyrosine kinases, ion channels, and downstream effectors^5,35,36^. Several studies have suggested that neuronal cilia influence brain function through signaling pathways mediated by these receptors. In the adult brain, G protein–coupled receptors, such as somatostatin receptor 3 and serotonin receptor 6, are preferentially localized to the ciliary membrane of neurons^37,38^. The role of hippocampal primary cilia in learning and memory, including contextual memory and novel object recognition, has been examined through deletion of ciliary proteins, such as somatostatin receptor 3^17^ and ACIII^33^. Although these proteins are enriched in neuronal cilia within the hippocampus, the functional significance of their ciliary localization and signaling, as well as the contribution of primary cilia to adult hippocampal function, remain incompletely understood. Because deletion of these ciliary proteins did not disrupt the formation of primary cilia, these proteins are probably essential for ciliary or cellular signaling rather than ciliogenesis. Therefore, behavioral deficits observed in these models may reflect disrupted signaling outside the cilium or other functions of these proteins.

The impact of primary cilia loss on neural function and behavior has been investigated in several studies, with mixed findings. Conditional deletion of *Ift88* in hippocampal and cortical neurons by using *Emx1^Cre^*, which deletes cilia during early development, impairs synaptic properties and behavioral deficits in aversive learning, memory, and novel object recognition^16^. Inducible global *Ift88* deletion in adult mice (*Ift88^fl/fl^; UBC^Cre^/ERT2*) impairs trace fear conditioning and Morris water maze performance and alters electroencephalographic activity^19^. In addition, AAV-mediated deletion or knockdown of ciliary genes in adult hippocampal neurons produces distinct effects on synaptic plasticity and memory^18,20^. Although these studies provide potential roles for primary cilia in adult brain function, whether selective loss of primary cilia in mature neurons directly alters neuronal physiology and behavior remained unresolved.

To minimize potential confounding effects from cilia-independent functions of ciliary proteins or ciliary loss in non-neuronal cell types, we selectively depleted primary cilia from mature hippocampal neurons. Therefore, we focused on neuron-specific cilia loss in the adult brain, rather than global deletion of cilia or ciliary proteins. Despite substantial loss of neuronal cilia in CA1 neurons and CA3 neurons, ciliary depletion did not alter neuronal excitability or synaptic plasticity. These findings were further supported by the lack of neuronal or synaptic deficits resulting from acute or virus-mediated cilia deletion in adult hippocampal neurons.

Several mechanisms may account for the absence of detectable physiological and behavioral phenotypes after primary cilia deletion in mature hippocampal neurons. One possibility is that primary cilia play a more prominent role in the development of neurons than in maintaining the physiological properties of mature neurons. Alternatively, signaling pathways outside the primary cilium may compensate for ciliary loss in adult hippocampus, or ciliary signaling may serve a modulatory (rather than essential) role under basal conditions. Consistent with the latter possibility, neither chronic genetic deletion nor acute or virus-mediated cilia ablation altered intrinsic excitability, synaptic transmission, or synaptic plasticity, indicating that mature hippocampal neurons maintain normal function in the absence of primary cilia. Although cilia depletion was incomplete in some hippocampal subfields and the subgranular zone neurogenic niche remained intact, these factors are unlikely to fully account for the lack of the phenotypes, given the robust cilia deletion achieved in CA1 and CA3 neurons. It should be noted that the absence of behavioral phenotypes may be attributable, at least in part, to the lower-than-expected performance observed in the behavioral assays, especially in *Grik4^Cre^* mouse strain; these negative findings should therefore be interpreted with this limitation in mind. Finally, our findings do not exclude important roles of primary cilia in the brain under pathologic conditions, stress, aging, or in response to altered neuronal activity, where ciliary signaling may become more prominent^39–46^.

Together, our results indicate that deletion of primary cilia from adult hippocampal neurons is not sufficient to produce detectable changes in neuronal function or behavior under the conditions examined. Rather than contradicting prior studies, these findings suggest that ciliary contributions to adult hippocampal function may be context-, circuit-, and cell type–dependent. Our results support a model in which primary cilia are dispensable for basal hippocampal function in mature excitatory neurons but may regulate neuronal signaling in specific cellular, physiological, or pathological conditions. Future studies combining systematic spatiotemporal manipulation of primary cilia with cell type– and brain region– specific approaches will be important for defining their precise roles in neural function and behavior.

## Methods and Materials

### Animals

Both male and female (3–5 months old) mice were used for all experiments. Mice with the floxed *Ift88* allele (JAX stock 022409) and the *Camk2a^Cre^* (JAX stock 005359), *Grik4^Cre^* (JAX stock 006474), and *Ai14* (JAX stock 007914) mouse strains were purchased from the Jackson Laboratory. The generation of *Arl13b-GFP*^30^ mouse line has been reported previously. All mouse strains were backcrossed onto the C57BL/6J genetic background for more than 5 generations. For most experiments, the experimenters were blinded to the genotype or treatment. The care and use of animals were reviewed and approved by the Institutional Animal Care and Use Committee at St. Jude Children’s Research Hospital.

### Immunohistochemistry and image analysis

Mice were deeply anesthetized by intraperitoneal injection of Avertin (0.2 ml/10 g, 1.25% v/v solution) and intracardially perfused with 4% paraformaldehyde in 0.1 mol/L phosphate-buffer solution (PBS) (pH 7.4). After perfusion, mice were rapidly decapitated using a rodent guillotine (World Precision Instruments), and brains were dissected and post-fixed overnight. Each brain was sliced (50-µm thickness) coronally with a vibratome (Leica VT1000S) post-fixed overnight. Each brain was sliced (50-µm thickness) coronally with a vibratome (Leica) and stored in PBS. Brain sections were incubated with blocking buffer (5% goat serum, 0.2% TritonX-100, 3% bovine serum albumin in PBS) for 1 h at room temperature. Next, the sections were incubated with primary antibodies against ACIII (Santa Cruz, sc-588, 1:250), red fluorescent protein (Rockland, 600-401-379, 1:500), or DCX (Abcam, ab18723; 1:1000) for 2 days at 4°C. Appropriate Alexa Fluor–conjugated secondary antibodies (Thermo Fisher Scientific, 1:1000) were used to detect primary antibody binding via 2-day incubation at 4°C. DAPI (Invitrogen) was used as a nuclear counterstain. All fluorescence images were acquired on a Zeiss780 confocal microscope. Cilia in the hippocampus were counted by reconstructing 3-dimensional z-stacked images of ACIII^+^ cells in the identified field.

### Viral infection and surgery

For *in vivo* viral injections, *Ift88^+/^* ^+^ mice and *Ift88^fl/fl^* mice were anesthetized with isoflurane (2% induction and 1.5% maintenance in pure oxygen). Mice were placed in a stereotaxic apparatus, and AAV5-hSyn-GFP-Cre vectors (UNC Vector Core) were injected into the dorsal hippocampus by using a 33-gauge metal cannula (Plastics One). The injection coordinates were AP: –1.8 mm, ML: ±1.0 mm, DV: –2.0 mm. Following injection, incisions were sutured, and mice were allowed to recover. Experiments were performed approximately 3 weeks after viral injection.

### Brain slice preparation and whole-cell electrophysiology

Transverse hippocampal slices (400-µm thickness) were prepared from adult mice, as previously described ^47,48^. Mice were rapidly decapitated using the rodent guillotine (World Precision Instruments), and brains were removed and placed in cold (4 °C) dissecting media containing (in mM) 125 choline-Cl, 2.5 KCl, 0.4 CaCl_2_, 6 MgCl_2_, 1.25 NaH_2_PO_4_, 26 NaHCO_3_, and 20 glucose (300–310 mOsm), equilibrated with 95% O_2_/5% CO_2_. Slices were made using a vibratome (Leica VT1200) and transferred to a recording chamber, where they were continuously superfused (2-3 mL/min) with artificial cerebrospinal fluid (ACSF) containing (in mM) 125 NaCl, 2.5 KCl, 2 CaCl_2_, 2 MgSO_4_, 1.25 NaH_2_PO_4_, 26 NaHCO_3_, and 10 glucose (approximately 295 mOsm), continuously bubbled with 95% O_2_/5% CO_2_ at 31-32°C. Whole-cell recordings were performed at 32°C by using either a potassium gluconate–based internal solution containing (in mM) 115 potassium gluconate, 20 KCl, 10 HEPES, 4 MgCl_2_, 0.1 EGTA, 4 MgATP, 0.4 NaGTP, and 10 sodium creatine phosphate (pH 7.4, 290-295 mOsm) for current-clamp recordings or a cesium-based internal solution containing (in mM) 125 CsMeSO_3_, 2 CsCl, 10 HEPES, 0.1 EGTA, 4 MgATP, 0.3 NaGTP, 10 sodium creatine phosphate, 5 QX-314, and 5 tetraethylammonium chloride, supplemented with 10-25 μM Alexa Fluor 594 (pH adjusted with CsOH, 290-295 mOsm) for voltage-clamp recordings. QX-314 was included to block AP generation. Intrinsic excitability was assessed by applying graded depolarizing current injections from the resting MP. Signals were recorded using a Multiclamp 700B amplifier, digitized with a Digidata 1550B interface at 20 kHz, filtered at 2 kHz, and acquired using Clampex 10.7 software (Axon Instruments). The liquid junction potential (−10 mV) was corrected offline.

For two-photon calcium imaging, intracellular calcium transients were measured by line-scanning the apical dendrites of CA1 pyramidal neurons during trains of back-propagating APs (20 APs at 100 Hz), before and after acute cilium ablation in the same neuron. Alexa Fluor 594 was included in all recordings to visualize neuronal morphology. Only morphologically intact CA1 pyramidal neurons with dendritic spines and a well-defined primary apical dendrite were included in the analysis.

In paired recordings, the presynaptic and postsynaptic neurons were filled with potassium gluconate–based internal solution. Acute ablation of the presynaptic primary cilium was performed by line-scanning across the base of the cilium using two-photon laser-scanning microscopy (Ultima imaging system, Bruker) equipped with a Chameleon Ultra femtosecond-pulsed laser (Coherent, 820 nm) and 60× water-immersion infrared objectives (Olympus). EPSPs were recorded before and after cilium ablation.

For LTP experiments, whole-cell EPSCs were recorded with the cesium-based internal solution from CA1 pyramidal neurons during Schaffer collateral stimulation. Following a stable baseline, LTP was induced using 200-Hz tetanic stimulation (40 stimulations, repeated 10 times, 5 s apart). EPSC amplitudes were normalized to the pre-tetanus baseline and monitored for 30–40 min after induction. The same method was used to assess LTP in hippocampal slices from *Ift88^fl/fl^* mice after AAV-GFP-Cre–mediated deletion of primary cilia.

### Hippocampal field potential recording

Field recordings were made using a submerged recording chamber setup (2 complete setups, 4 chambers each; Campden Instruments), Digidata digitizers and computer-controlled microelectrode amplifiers (Molecular Devices), STG2004 stimulators (Multichannel Systems), and pClamp software (Molecular Devices), with each chamber recorded independently. Field excitatory postsynaptic potentials (fEPSPs) were recorded from the CA1 stratum radiatum by using an extracellular glass pipette (3-5 MΩ) filled with ACSF. Schaffer collateral fibers were stimulated with a bipolar tungsten electrode (FHC) placed 200-300 μm from the recording electrode. Input/output curves were first generated for each slice, and the stimulus intensity was adjusted to evoke approximately one-third of the maximal fEPSP response. Paired-pulse facilitation was then assessed using interstimulus intervals of 20, 50, 100, 200, and 1000 ms. For the LTP test, paired stimuli (40-ms interval) were delivered for at least 20 min to establish a stable baseline before LTP was induced with three rounds of 200-Hz tetanic stimulation. Each round was separated by 5 min and consisted of 10 trains of 200-Hz stimulation (200-ms duration; 40 pulses) delivered every 5 s. Paired post-tetanic responses (50-ms interstimulus interval) were recorded every 30 s for 90 min.

### Fear conditioning

Contextual and auditory cued fear conditioning were tested using a conditioning chamber (VFC-008 and NIR-022 MD, Med Associate) housed within a sound-attenuating cabinet as previously described^49^. Behavior was recorded using a camera and analyzed with VideoFreeze Software (v.3.02.00.00). All behavioral apparatuses were cleaned with 70% ethanol between trials. Training (Day 1): mice were placed in the conditioning chamber with the white house light on and allowed to explore the testing chamber for 5 min. Each conditioned stimulus (CS)–unconditioned stimulus (US) pairing consisted of 30-s auditory tone (10 kHz, 75 dB SPL), followed by a 0.5-mA foot shock during the final 2 s of the tone. After a 1-min recovery period, seven additional CS–US pairings were delivered. Mice remained in the chamber for 2 min after the final pairing before being returned to their home cage. Contextual test (Day 2): approximately 24 h after training, mice were returned to the conditioning chamber, and freezing behavior was recorded for 5 min. Cued test (Day 3): mice were placed in a modified chamber with novel walls and flooring and allowed to explore for 2 min. The auditory CS (30 s) was then presented without foot shock, followed by a 30-s recovery period. This sequence was repeated four additional times.

### Novel object recognition

Novel object recognition was tested in an open-field arena as previously described^50^, and behavior was recorded and analyzed using TopScan software (v.3.2, CleverSys Inc.). During the familiarization phase, two identical objects were placed approximately 6 cm from opposite walls of the arena. Mice were placed in the arena facing away from the objects and allowed to explore freely for 4 min before being returned to their home cages. At 24 h later, one familiar object was replaced with a novel object in the same location. Mice were again placed in the arena facing away from the objects and allowed to explore for 5 min. Recognition memory was evaluated by calculating object preference as follows: familiar object preference (%) = (time exploring the familiar object/total object exploration time) × 100; novel object preference (%) = (time exploring the novel object/total object exploration time) × 100.

### Morris water maze

Spatial learning and memory were tested using the Morris water maze, as previously described^51^. A circular pool was filled with opaque water and contained a hidden platform submerged 2 cm below the water surface. Mice were allowed to habituate to the testing room for 1 h before the experiment was started. Swim paths were recorded and analyzed using the TopScan video-tracking system. During the acquisition phase, mice were trained to locate the hidden platform in four trials per day with a 1-min intertrial interval for 5 consecutive days. Starting positions were counterbalanced daily using a Latin square design. Escape latency (time to reach the hidden platform) was used to measure spatial learning. Probe trials were performed 1 h and 72 h after completion of the final training session. During each 1-min probe trial, the platform was removed, and the mice were released from the starting position farthest from the former platform location. Time spent in the target quadrant was used as the measure of spatial memory retrieval. To avoid hypothermia, mice were dried with paper towels and placed in warm holding cages after each training and testing session.

### Open-field test

Anxiety-like behavior was assessed using the open-field test. The test apparatus consisted of a 16 × 16-in arena with 15-in high walls, overhead lighting, and a camera for automated video tracking and analysis (TopScan). Mice were acclimated to the testing room for 1 h on 2 consecutive days. On the third day, mice were placed in the arena and allowed to explore freely for 10 min.

### Three-chamber test

Sociability and social recognition were assessed using a three-chamber apparatus. Mice were initially placed in the center chamber with empty containers positioned in the two side chambers and allowed to habituate for 10 min. For the sociability test, the doors to the side chambers were opened, and an age- and sex-matched stranger mouse was placed in one container while the other remained empty. The test mouse was then allowed to freely explore all three chambers for 10 min. For the social recognition test, a second unfamiliar age- and sex-matched mouse was placed in the previously empty container 1h later, and the test mouse was allowed to explore all three chambers for an additional 10 min. Behavior was recorded and analyzed using TopScan software.

### Rotarod test

Motor coordination and performance were assessed using the rotarod test. Mice were placed on a rotating rod (3.18-cm diameter) separated into 11.4-cm wide lanes (ROTOR-ROD™ System). The rotation speed was gradually increased from 0 to 20 rpm over 4 min and then maintained until the mouse fell. Mice were tested in two training blocks, each consisting of two trials per day for 5 consecutive days, separated by a 9-day interval. Latency to fall (s) and distance travelled (cm) were recorded.

### Statistical analyses

Statistical comparisons were performed using two-tailed Student’s *t*-test or Welch’s unpaired *t*-test for two groups and one-way or two-way ANOVA for multiple groups, followed by Holm–Sidak’s pairwise comparisons when the normality assumption was met. All statistical tests were performed using an alpha level of 0.05. *F*-values were reported for ANOVA. All analyses were computed using Prism 10 (GraphPad software) or Sigma Plot (Systat software). Specific statistical details are provided in the figure legends.

## Acknowledgments

We thank the members of the Zakharenko lab for providing comments and Angela McArthur for editing the manuscript. We thank David Clapham for providing Arl13b-GFP mice. Research reported in this publication was supported by the National Institute on Deafness and Other Communication Disorders of the National Institutes of Health under Award Number R01DC021511. This work was also supported by the American Lebanese Syrian Associated Charities (ALSAC), the BBRF Young Investigator Award (T.- Y.E.), and the BBRF Distinguished Investigator Award (SSZ). The funding sources had no role in the study design, data collection, data analysis, decision to publish, or preparation of the manuscript.

## Author contributions

T.-Y. E. and S.S.Z. designed the study. T.-Y. E. performed behavioral and immunohistochemical studies, viral injection experiments, and maintained all mouse lines. I.T.B. and B. J.W.T performed the electrophysiological assays. B.J.W.T and D. E. performed behavioral experiments. S.S.Z. provided reagents and equipment. T.-Y. E. wrote the original draft, and all authors reviewed the manuscript.

## Data availability

All relevant data associated with the published study are present in the paper or the Supplementary Information. Additional data relating to this paper are available upon request from the corresponding author.

## Ethics declarations

This study was approved by the Institutional Animal Care and Use Committee at St. Jude Children’s Research Hospital (Protocol no: 3177). All procedures were carried out in strict accordance with the ARRIVE guidelines and policies of the Institutional Animal Care and Use Committee at St. Jude Children’s Research Hospital.

## References

1. Youn, Y. H. & Han, Y.-G. Primary cilia in brain development and diseases. Am. J. Pathol. 188, 11–22 (2018).

2. Huangfu, D. et al. Hedgehog signalling in the mouse requires intraflagellar transport proteins. Nature 426, 83–87 (2003).

3. Ansley, S. J. et al. Basal body dysfunction is a likely cause of pleiotropic Bardet–Biedl syndrome. Nature 425, 628–633 (2003).

4. Guemez-Gamboa, A., Coufal, N. G. & Gleeson, J. G. Primary cilia in the developing and mature brain. Neuron 82, 511–521 (2014).

5. Mill, P., Christensen, S. T. & Pedersen, L. B. Primary cilia as dynamic and diverse signalling hubs in development and disease. Nat. Rev. Genet. 24, 421–441 (2023).

6. Pedersen, L. B. & Rosenbaum, J. L. Intraflagellar Transport (IFT). in Current Topics in Developmental Biology (ed. Bradley K. Yoder) vol. 85 23–61 (University of Copenhagen, Yale University, 2008).

7. Goetz, S. C. & Anderson, K. V. The primary cilium: a signalling centre during vertebrate development. Nat. Rev. Genet. 11, 331–344 (2010).

8. Berbari, N. F., O’Connor, A. K., Haycraft, C. J. & Yoder, B. K. The primary cilium as a complex signaling center. Current Biology 19, R526–R535 (2009).

9. Lancaster, M. A. & Gleeson, J. G. The primary cilium as a cellular signaling center: lessons from disease. Curr. Opin. Genet. Dev. 19, 220–229 (2009).

10. Hildebrandt, F., Benzing, T. & Katsanis, N. Ciliopathies. New England Journal of Medicine 364, 1533–1543 (2011).

11. Braun, D. A. & Hildebrandt, F. Ciliopathies. Cold Spring Harb. Perspect. Biol. 9, a028191 (2017).

12. Kumamoto, N. et al. A role for primary cilia in glutamatergic synaptic integration of adult-born neurons. Nat. Neurosci. 15, 399–405 (2012).

13. Chizhikov, V. V. et al. Cilia proteins control cerebellar morphogenesis by promoting expansion of the granule progenitor pool. The Journal of Neuroscience 27, 9780–9789 (2007).

14. Higginbotham, H. et al. Arl13b in primary cilia regulates the migration and placement of interneurons in the developing cerebral cortex. Dev. Cell 23, 925–938 (2012).

15. Tong, C. K. et al. Primary cilia are required in a unique subpopulation of neural progenitors. Proceedings of the National Academy of Sciences 111, 12438–12443 (2014).

16. Berbari, N. F. et al. Hippocampal and cortical primary cilia are required for aversive memory in mice. PLoS One 9, e106576 (2014).

17. Einstein, E. B. et al. Somatostatin signaling in neuronal cilia is critical for object recognition memory. The Journal of Neuroscience 30, 4306–4314 (2010).

18. Jovasevic, V. et al. Primary cilia are required for the persistence of memory and stabilization of perineuronal nets. iScience 24, 102617 (2021).

19. Strobel, M. R., Zhou, Y., Qiu, L., Hofer, A. M. & Chen, X. Temporal ablation of the ciliary protein IFT88 alters normal brainwave patterns. Sci. Rep. 15, 347 (2025).

20. Rhee, S., Kirschen, G. W., Gu, Y. & Ge, S. Depletion of primary cilia from mature dentate granule cells impairs hippocampus-dependent contextual memory. Sci. Rep. 6, 34370 (2016).

21. Haycraft, C. J. et al. Intraflagellar transport is essential for endochondral bone formation. Development 134, 307–316 (2007).

22. Tsien, J. Z. et al. Subregion- and cell type-restricted gene knockout in mouse brain. Cell 87, 1317–1326 (1996).

23. Sperow, M. et al. Phosphatase and tensin homologue (PTEN) regulates synaptic plasticity independently of its effect on neuronal morphology and migration. Journal of Physiology 590, (2012).

24. Zakharenko, S. S. et al. Presynaptic BDNF required for a presynaptic but not postsynaptic component of LTP at hippocampal CA1-CA3 synapses. Neuron 39, (2003).

25. Nakazawa, K. et al. Requirement for Hippocampal CA3 NMDA receptors in associative memory recall. Science (1979). 297, 211–218 (2002).

26. Martin, S. J., Grimwood, P. D. & Morris, R. G. Synaptic plasticity and memory: an evaluation of the hypothesis. Annu Rev Neurosci 23, 649–711 (2000).

27. Zakharenko, S. S., Zablow, L. & Siegelbaum, S. A. Visualization of changes in presynaptic function during long-term synaptic plasticity. Nat. Neurosci. 4, (2001).

28. Milner, B., Squire, L. R. & Kandel, E. R. Cognitive neuroscience and the study of memory. Neuron 20, 445–468 (1998).

29. Nakazawa, K. et al. Requirement for hippocampal CA3 NMDA receptors in associative memory recall. Science (1979). 297, 211–218 (2002).

30. Delling, M., DeCaen, P. G., Doerner, J. F., Febvay, S. & Clapham, D. E. Primary cilia are specialized calcium signalling organelles. Nature 504, 311–314 (2013).

31. Sabatini, B. L., Oertner, T. G. & Svoboda, K. The life cycle of Ca(2+) ions in dendritic spines. Neuron 33, 439–452 (2002).

32. Richardson, R. J., Blundon, J. A., Bayazitov, I. T. & Zakharenko, S. S. Connectivity patterns revealed by mapping of active inputs on dendrites of thalamorecipient neurons in the auditory cortex. Journal of Neuroscience 29, 6406–6417 (2009).

33. Wang, Z., Phan, T. & Storm, D. R. The type 3 adenylyl cyclase is required for novel object learning and extinction of contextual memory: role of cAMP signaling in primary cilia. The Journal of Neuroscience 31, 5557–5561 (2011).

34. Tereshko, L., Turrigiano, G. G. & Sengupta, P. Primary cilia in the postnatal brain: Subcellular compartments for organizing neuromodulatory signaling. Curr. Opin. Neurobiol. 74, 102533 (2022).

35. Christensen, S. T., Morthorst, S. K., Mogensen, J. B. & Pedersen, L. B. Primary Cilia and coordination of receptor tyrosine kinase (RTK) and transforming growth factor β (TGF-β) signaling. Cold Spring Harb. Perspect. Biol. 9, a028167 (2017).

36. Hilgendorf, K. I., Johnson, C. T. & Jackson, P. K. The primary cilium as a cellular receiver: organizing ciliary GPCR signaling. Curr. Opin. Cell Biol. 39, 84–92 (2016).

37. Brailov, I. et al. Localization of 5-HT6 receptors at the plasma membrane of neuronal cilia in the rat brain. Brain Res. 872, 271–275 (2000).

38. Domire, J. S. & Mykytyn, K. Markers for neuronal cilia. in 111–121 (2009). doi:10.1016/S0091-679X(08)91006-2.

39. Pelaz, S. G. et al. Amygdala astrocyte primary cilium mechanisms contribute to stress behaviours. Nature 10.1038/s41586-026-10874-0 (2026) doi:10.1038/s41586-026-10874-0.

40. Sheu, S.-H. et al. A serotonergic axon-cilium synapse drives nuclear signaling to alter chromatin accessibility. Cell 185, 3390–3407.e18 (2022).

41. Zhai, D. et al. The role of primary cilia in physiological and pathological states of the central nervous system. Journal of Genetics and Genomics 53, 577–593 (2026).

42. Fitzsimons, L. A. et al. The nociceptor primary cilium contributes to mechanical nociceptive threshold and inflammatory and neuropathic pain. The Journal of Neuroscience 44, e1265242024 (2024).

43. Ma, R., Kutchy, N. A., Chen, L., Meigs, D. D. & Hu, G. Primary cilia and ciliary signaling pathways in aging and age-related brain disorders. Neurobiol. Dis. 163, 105607 (2022).

44. Alhassen, W. et al. Patterns of cilia gene dysregulations in major psychiatric disorders. Prog. Neuropsychopharmacol. Biol. Psychiatry 109, 110255 (2021).

45. Baek, H. et al. Primary cilia modulate TLR4-mediated inflammatory responses in hippocampal neurons. J. Neuroinflammation 14, 189 (2017).

46. Yang, J. et al. Primary ciliary protein kinase A activity in the prefrontal cortex modulates stress in mice. Neuron 113, 1276–1289.e5 (2025).

47. Bayazitov, I. T., Richardson, R. J., Fricke, R. G. & Zakharenko, S. S. Slow presynaptic and fast postsynaptic components of compound long-term potentiation. Journal of Neuroscience 27, (2007).

48. Zakharenko, S. S., Zablow, L. & Siegelbaum, S. A. Altered presynaptic vesicle release and cycling during mGluR-dependent LTD. Neuron 35, (2002).

49. Eom, T. Y., Bayazitov, I. T., Anderson, K., Yu, J. & Zakharenko, S. S. Schizophrenia-related microdeletion impairs emotional memory through microRNA-dependent disruption of thalamic inputs to the amygdala. Cell Rep. 19, 1532–1544 (2017).

50. Devaraju, P. et al. Haploinsufficiency of the 22q11.2 microdeletion gene Mrpl40 disrupts short-term synaptic plasticity and working memory through dysregulation of mitochondrial calcium. Mol Psychiatry 22, 1313–1326 (2017).

51. Earls, L. R. et al. Dysregulation of presynaptic calcium and synaptic plasticity in a mouse model of 22q11 deletion syndrome. J Neurosci 30, 15843–15855 (2010).

